# Ancestral role of Pax6 in chordate brain regionalization

**DOI:** 10.1101/2024.05.03.592360

**Authors:** Zbynek Kozmik, Iryna Kozmikova

## Abstract

The *Pax6* gene is essential for eye and brain development across various animal species. Here, we investigate the function of *Pax6* in the development of the anterior central nervous system (CNS) of the invertebrate chordate amphioxus using CRISPR/Cas9-induced genome editing. Specifically, we examined Pax6 mutants featuring a 6bp deletion encompassing two invariant amino acids in the conserved paired domain, hypothesized to impair Pax6 DNA-binding capacity and gene regulatory functions. Although this mutation did not result in gross morphological changes in amphioxus larvae, it demonstrated a reduced ability to activate Pax6-responsive reporter gene, suggesting a hypomorphic effect. Expression analysis in mutant larvae revealed changes in gene expression within the anterior CNS, supporting the conserved role of *Pax6* gene in brain regionalization across chordates. Additionally, our findings lend support to the hypothesis of a zona limitans intrathalamica (ZLI)-like region in amphioxus, suggesting evolutionary continuity in brain patterning mechanisms.

## INTRODUCTION

*Pax6* is a member of the homeobox gene family, which also contains a DNA-binding paired-box motif originally identified in *Drosophila* (*1*). The paired domains of Pax6 proteins exhibit a high degree of sequence conservation; vertebrate Pax6 proteins display nearly identical paired domains, whereas invertebrate Pax6 proteins show more than 90% sequence homology with their mouse Pax6 (*2*). Since its discovery in 1991 (*3*), studies on *Pax6* lead to the transformative thinking regarding the genetic programs orchestrating eye morphogenesis as well as the origin and evolution of diverse visual systems. The uncovering of the *Pax6* gene as an essential factor in eye development within both mice (*4, 5*) and *Drosophila* (*6, 7*) has given rise to the concept of a "master control gene for eye morphogenesis and evolution," alongside the hypothesis of a monophyletic origin of eyes in metazoans (*8*). This idea presented a stark contrast to the perspective originally posited by Salvini-Plawen and Mayr, which suggested a diverse, independent origin of photoreceptor organs across numerous species (*9*). The theory about a ‘master control gene,’ has propelled a wave of scientific investigation into the expression and function of Pax6 across diverse animal species (*10*).

Pax6 is a typical pleiotropic transcription factor that has been implicated in diverse biological processes, and it is known to regulate expression of a broad range of molecules, including transcription factors, cell adhesion and cell signaling molecules, hormones, and structural proteins (reviewed in (*11, 12*)). *Pax6* function is not restricted to the visual system as it is also essential for the development of the central nervous system and endocrine glands of vertebrates and invertebrates. The expression patterns of *Pax6* in the developing nervous systems of vertebrates, eyes included, show significant similarity (*5, 13–16*).

Heterozygous mice carrying the Small eye (Sey) *Pax6* gene mutation (*4*), which involves a premature stop codon, display a range of eye deficits including aniridia, which is a condition also observed in humans, as well as lens size (*17, 18*). These mice also exhibit abnormalities in the telencephalon, diencephalon, and metencephalon (*19*). Homozygous Sey mutants are not viable; the embryos exhibit profound brain and olfactory malformations (*5*). *Pax6* plays a role in establishing boundaries between regions of the central nervous system in the anteroposterior axis, at least in part, due to the regulation of homeobox-containing genes such as *En1*, *Pax2*, and *Lhx1* (*20–22*). The boundary between cortical and striatal regions of the telencephalon is dramatically altered in Sey mutants: radial glial fascicles do not form at the border, and the normal expression of R-cadherin and tenascin-C at the border is lost suggesting that *Pax6* regulates boundary formation between developing forebrain regions (*23*). Paired domain is necessary for the regulation of neurogenesis, cell proliferation and patterning effects of Pax6, since these aspects are severely affected in the developing forebrain of the Pax6Aey18 mice with a deletion in the PD but intact homeodomain and transactivation domain (*24*).

In *Xenopus*, mutations that result in truncated Pax6 proteins affect forebrain regionalization but do not completely eliminate eyes; rather, they lead to the formation of eye-like structures without lenses (*25*). It is hypothesized that an additional *Pax6.2* gene may compensate for these phenotypic alterations. In medaka, mutations in the individual *Pax6.1* or *Pax6.2* genes do not completely eliminate eyes either (*14, 26*). Despite the shift away from the "master control gene" concept, *Pax6* central role in eye and brain development is undeniable, continuing to make it an intriguing subject for evolutionary studies.

The cephalochordate amphioxus, owing to its unique phylogenetic position as the presumed closest living relative to the common ancestor of chordates, serves as a pivotal model for exploring chordate evolution and vertebrate innovations. In amphioxus, *Pax6* expression initiates during neurulation in the surface anterior ectoderm and neural plate. As neurulation progresses, strong expression becomes localized in the anterior ectoderm and the developing cerebral vesicle, the presumptive homolog of the vertebrate brain (*27*). Previous studies, utilizing electron microscopy and gene expression analysis, have suggested that the brain’s anterior part corresponding to the frontal eye, may represent a potential homolog of the vertebrate paired eyes (*28–30*). More recently, in light of new experimental evidence (*31*) it has been proposed that the molecular signature of the frontal eye exhibits similarities to both the vertebrate retina and hypothalamus (*32*). Our research investigates the impact of a mutation in the most conserved region of the *Pax6* gene on the anterior central nervous system of amphioxus.

## RESULTS

### Targeted mutagenesis of *Branchiostoma floridae Pax6* gene

To determine the functional role of *Pax6* gene in amphioxus central nervous system development we analyzed mutants generated by CRISPR/Cas9 genome editing. Targeting 5’end of the exon encoding the N-terminal half of paired domain produced an allele carrying a 6bp deletion (Fig. 1A) that was transmitted to F1 generation, and was designated Pax6ΔQL. Progeny of genetic crosses between Pax6ΔQL F1 (and F2) animals was genotyped to identify wild type, heterozygote, and homozygote embryos (Fig. 1B). No significant morphological changes were observed in the homozygote mutant amphioxus larvae at 4 days of development. The 6bp deletion results in the elimination of the two evolutionarily conserved amino acids found in both bilaterian and cnidarian Pax proteins. Our analysis (Fig. 1C) has shown that the respective QL amino acids are conserved in all nine human paralogues (PAX1-PAX9), in *Drosophila* Pax6, and even in the cnidarian PaxB that was previously characterized as a structural hybrid between a typical bilaterian Pax6 and Pax2 (*33*).

**Fig. 1.**
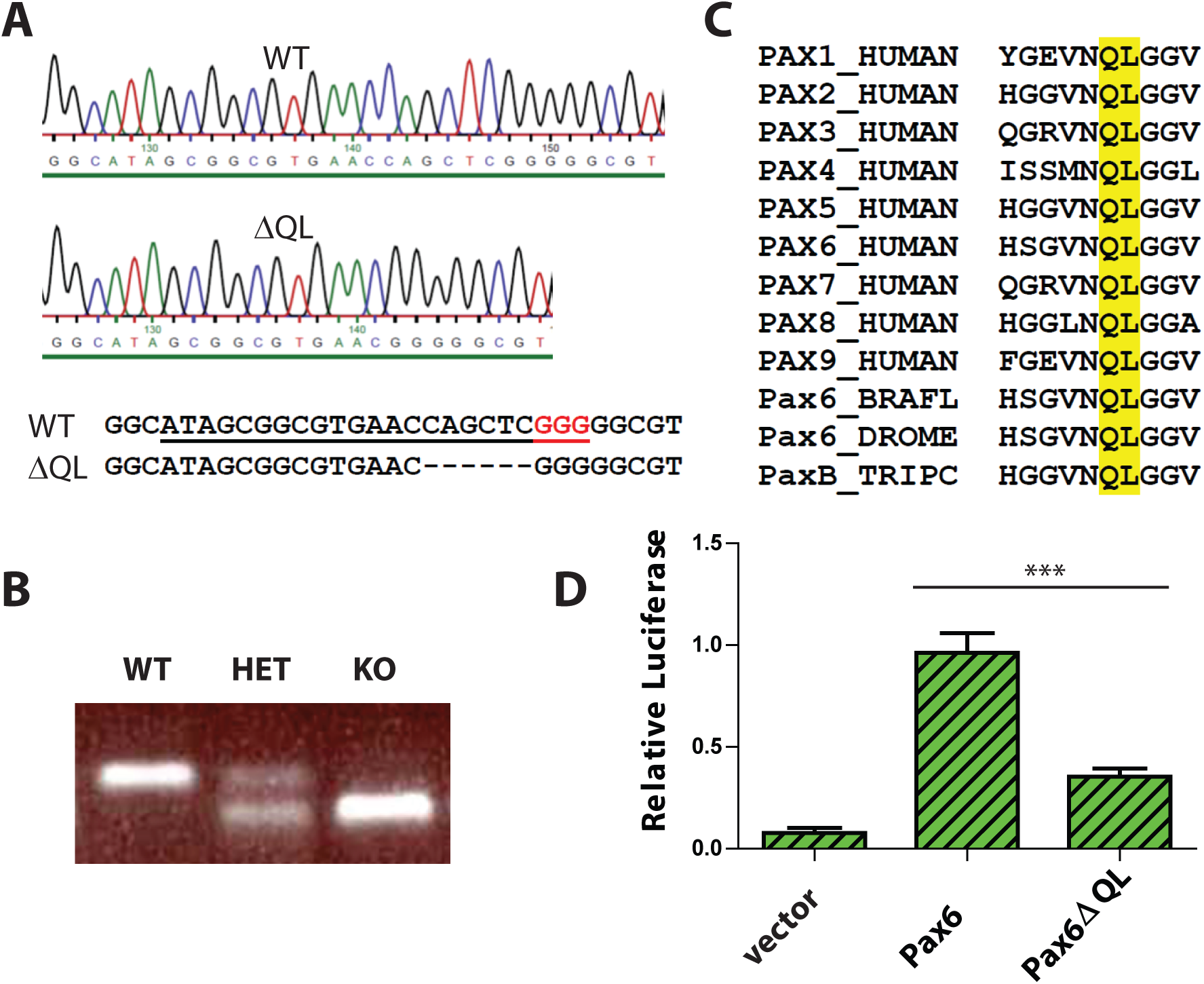
Targeted mutagenesis of *Branchiostoma floridae* Pax6 gene and characterization of the Pax6ΔQL mutant protein. (A) DNA sequencing results validating the 6bp deletion in the *Branchiostoma floridae* Pax6 gene. sgRNA used for CRISPR/Cas9 genome editing is underlined and PAM sequence highlighted red. (B) A representative PCR based genotyping of amphioxus embryos carrying wild type and Pax6ΔQL alleles (C) Amino acid sequence alignment of paired domain region for selected Pax genes. Amino acids QL deleted in mutant allele of *Branchiostoma floridae* Pax6 described here are highlighted by the yellow box. BRAFL, *Branchiostoma floridae*; DROME, *Drosophila melanogaster*; TRIPC, *Tripedalia cystophora*. (D) Reporter gene assays were performed by co-transfecting the indicated Pax6-encoding expression vector or empty expression vector with Pax6-responsive promoter construct. Triplicate assays were performed to obtain standard deviations and transfection efficiency was normalized by co-transfection of β-galactosidase expression plasmid.

We reasoned that elimination of two invariant amino acids of paired domain might compromise DNA binding ability of Pax6 and as a result diminish its ability to regulate target genes. This notion was further supported by the published structure of the Pax6 paired domain– DNA complex showing DNA contacts of the mutagenized amino acids with the phosphate backbone (*34*). To test the hypothesis we performed luciferase reporter gene assays using either wild type Pax6 or mutant Pax6ΔQL. Reporter gene assay revealed a strongly reduced capacity of the mutated protein to activate the Pax6 responsive promoter (Fig.1D). However, the observed residual activity of Pax6ΔQL as compared to the empty expression vector (Fig.1C) strongly suggests that the mutant allele generated by us here using CRISPR/Cas9 genome editing is hypomorhic.

### *Pax6* mutation affects the molecular organization and regionalization in the brain of amphioxus larvae

The results demonstrating the reduced activity of the mutated Pax6 protein encouraged us to closely examine the expression of marker genes in the region referred to as the frontal eye by Pergner et al.(*29*), or as the retina and hypothalamus according to Lacalli (*32*), as well as the proto-tectum and primary motor center, or dien-mesencephalon (suggested counterpart of vertebrate thalamus, pretectum, and midbrain). Up to now, it is not completely clear which concept should prevail, and for clarity, we will maintain the terminology proposed by Pergner et al.(*29*).

We analyzed the expression of *Six3/6*, *Otx*, and *Brn3* (Fig.2), which are found in photoreceptors of wild type larva (Fig. 2A-as3; Fig. 2Bd and bd; Fig. 2C-Cs1’; and c-cs1’). These genes showed no significant change in expression in these cells (Fig.4), indicating that the photoreceptors were likely unaffected. However, we did notice alterations in the expression of *Six3/6*, *Brn1/2/4*, *Lhx3*, and *Pax6* in other regions of the frontal eye. Specifically, *Six3/6* expression, seen at the boundary of the frontal eye with the proto-tectum in wild type larvae, was reduced in this region (Fig. 2A-A’ and 2a-a’; Fig. 2As3 and 2as3; Fig.4). Conversely, *Brn1/2/4* expression extended into the anterior frontal eye, affecting Row3, Row2, and even the photoreceptors (Fig. 2D-ds2). Lhx3 expression was significantly diminished in Row4 but not in Row3 cells, and *Pax6* showed reduced expression in Row4 and at the boundary of the frontal eye with the proto-tectum (Fig. 3B-bs2 and Fig. 4). Notably, there were no marked changes in the expression of *Otp* and *Lhx1* in Row4 (Fig. 3C-cs2 and 3D-ds2). However, both these genes, along with *Brn3*, were downregulated in the proto-tectum (Fig. 3C-cs2; Fig. 3D’-Dd and d’-dd; Fig. 4). Additionally, *Brn3* and *Lhx1* expression was slightly elevated at the boundary of the frontal eye with the proto-tectum (Fig. 2C-C’ and c-c’; Fig. 3Ds1-Ds2 and ds1-ds2; Fig. 4). Conversely, *Pax6* and *Otx* expression was reduced in this area (Fig. 2A-as3; Fig. 2B-bd; Fig. 3B-bs2; Fig.4). Apart from *Otp*, all genes expressed in the primary motor center, including *Pax6*, *Lhx3*, *Brn3*, and *Six3/6*, were downregulated (Fig.2A-as2; Fig. 2C-cs2; Fig. 3A-as2; Fig. 3B-bs2; Fig.4). In summary, our data suggest that the most significant changes due to the *Pax6* mutation occur in the posterior frontal eye, proto-tectum, and primary motor center (Fig.5A).

**Fig.2.**
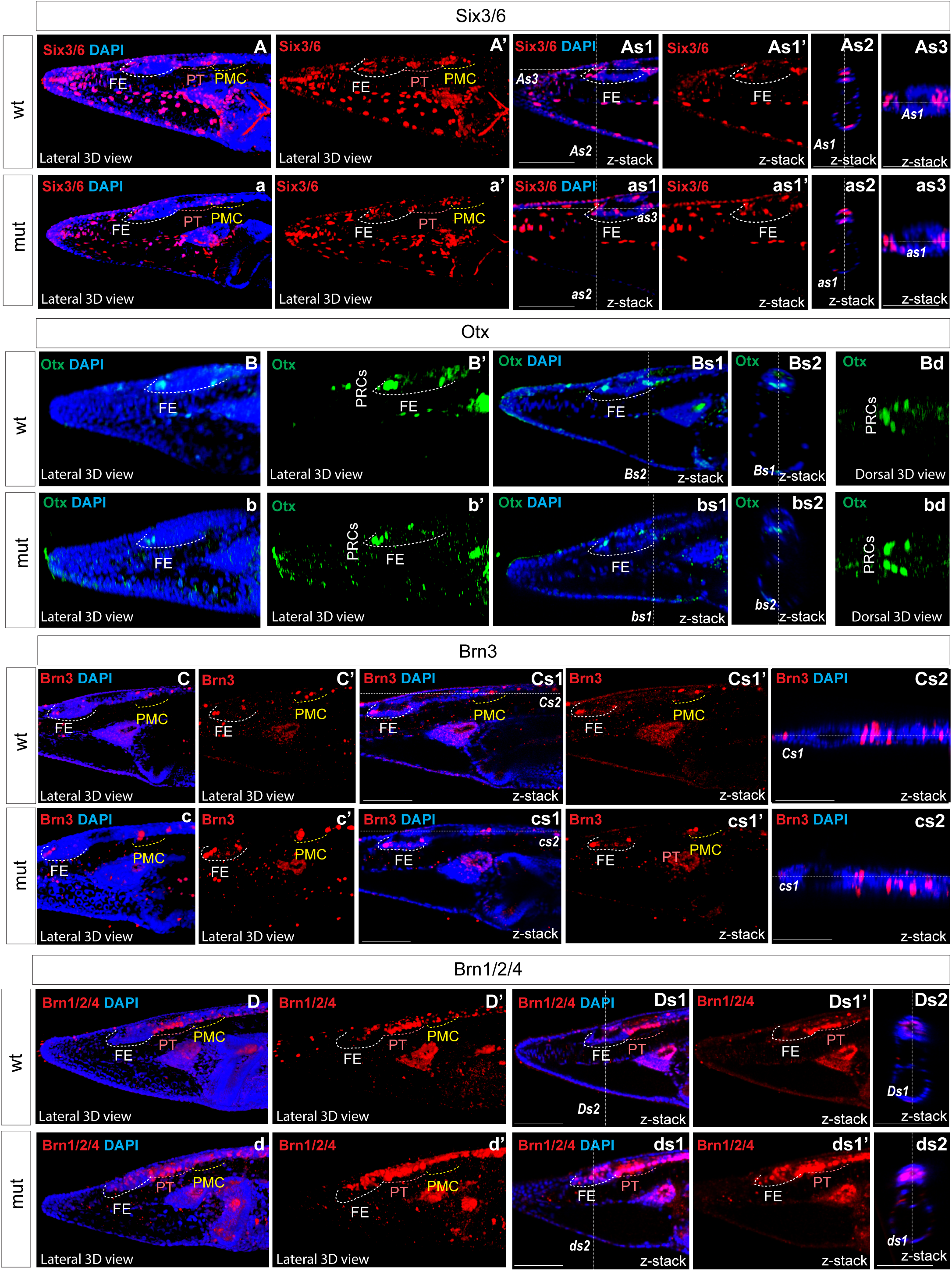
Expression of individual genes in amphioxus wild type (A-As3; B-Bs2; C-Cs2; D-Ds2) and Pax6ΔQL mutant embryos (a-as3; b-bs2; c-cs2; d-ds2). Wt-wild type embryos. Mut – mutant embryos FE – frontal eye. PT – proto-tectum PMC – primary motor center. PRCs – photoreceptors. The positions of individual z-slices (As2, As1, As3, as1, as2, as3, Bs1, Bs2, bs1, bs2 Cs1, Cs2, cs1, cs2, Ds1, Ds2, ds1, ds2) from complete z-stacks are indicated with dashed lines (As2, As1, As3, as1, as2, as3, Bs1, Bs2, bs1, bs2 Cs1, Cs2, cs1, cs2, Ds1, Ds2, ds1, ds2).

**Fig.3.**
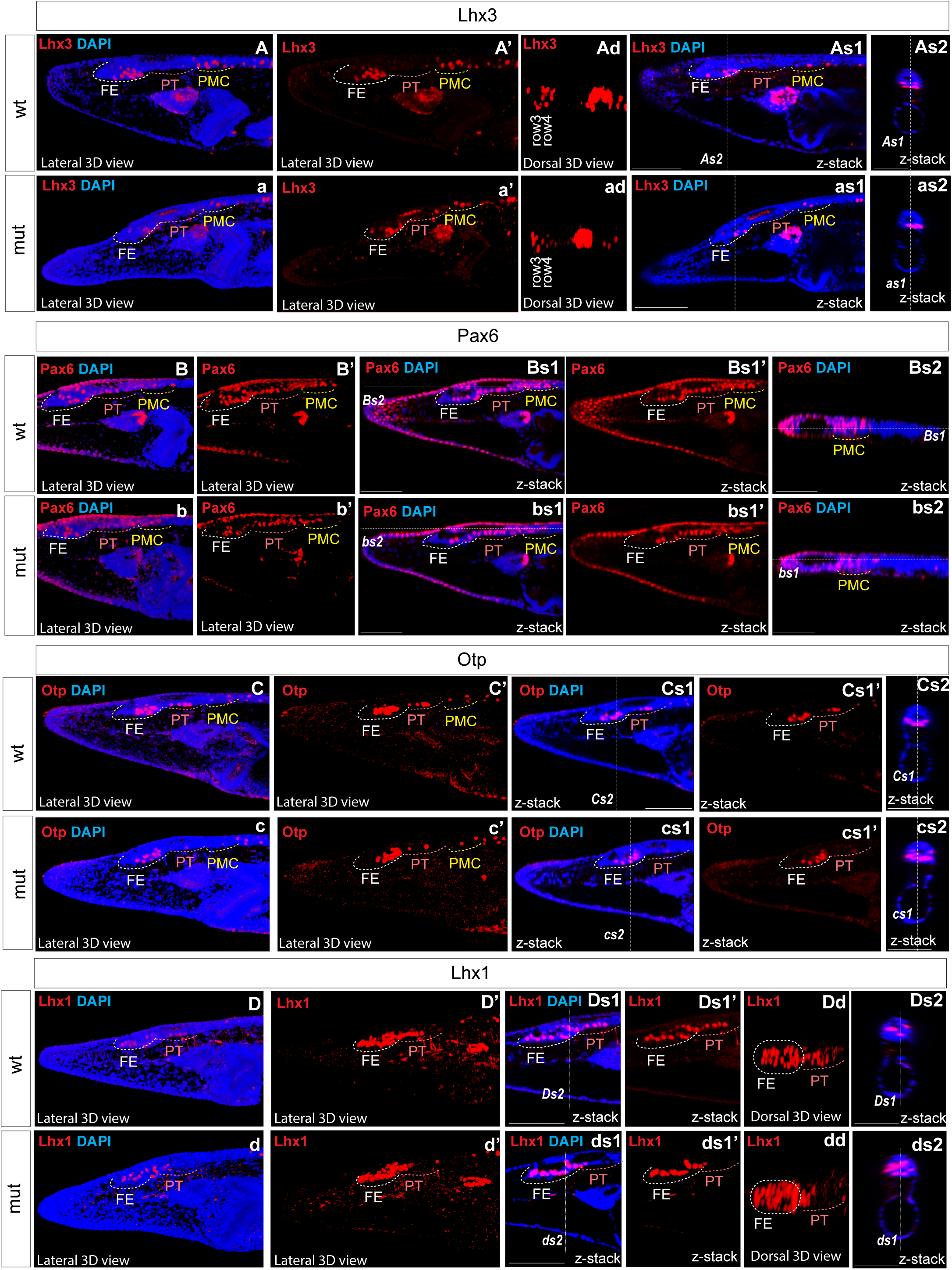
Expression of individual genes in amphioxus wild type (A-As3; B-Bs2; C-Cs2; D-Ds2) and Pax6ΔQL mutant embryos (a-as3; b-bs2; c-cs2; d-ds2). Wt-wild type embryos. Mut – mutant embryos FE – frontal eye. PT – proto-tectum PMC – primary motor center. PRCs – photoreceptors. The positions of individual z-slices (As2, As1, As3, as1, as2, as3, Bs1, Bs2, bs1, bs2 Cs1, Cs2, cs1, cs2, Ds1, Ds2, ds1, ds2) from complete z-stacks are indicated with dashed lines (As2, As1, As3, as1, as2, as3, Bs1, Bs2, bs1, bs2 Cs1, Cs2, cs1, cs2, Ds1, Ds2, ds1, ds2).

**Fig.4.**
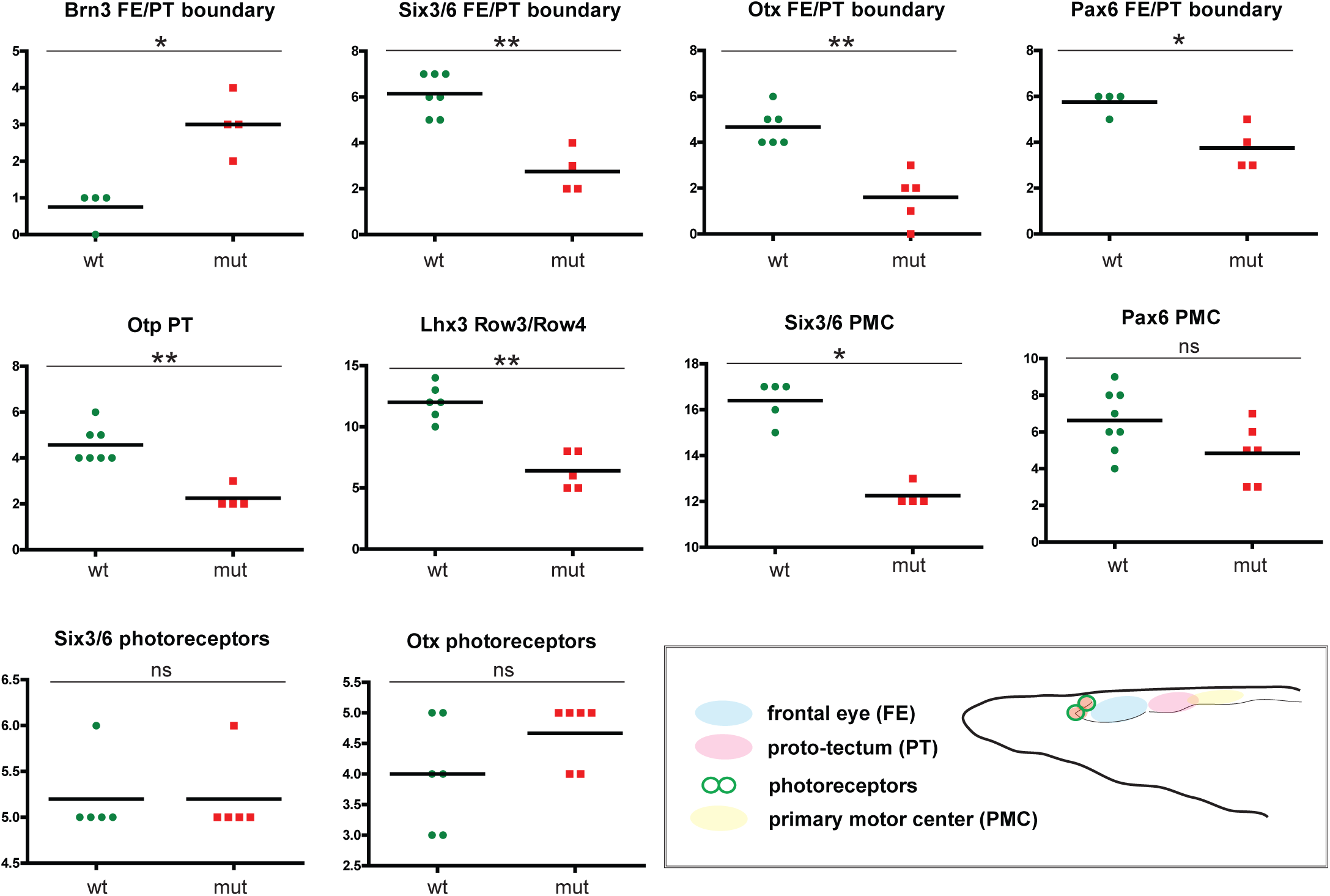
Difference in cell numbers between wild type and Pax6ΔQL mutant animals for the indicated regional markers. Mann-Whitney two tailed test was used for calculation of statistical significance. FE - frontal eye; PT-proto-tectum; PMC-primary motor center.

**Fig.5.**
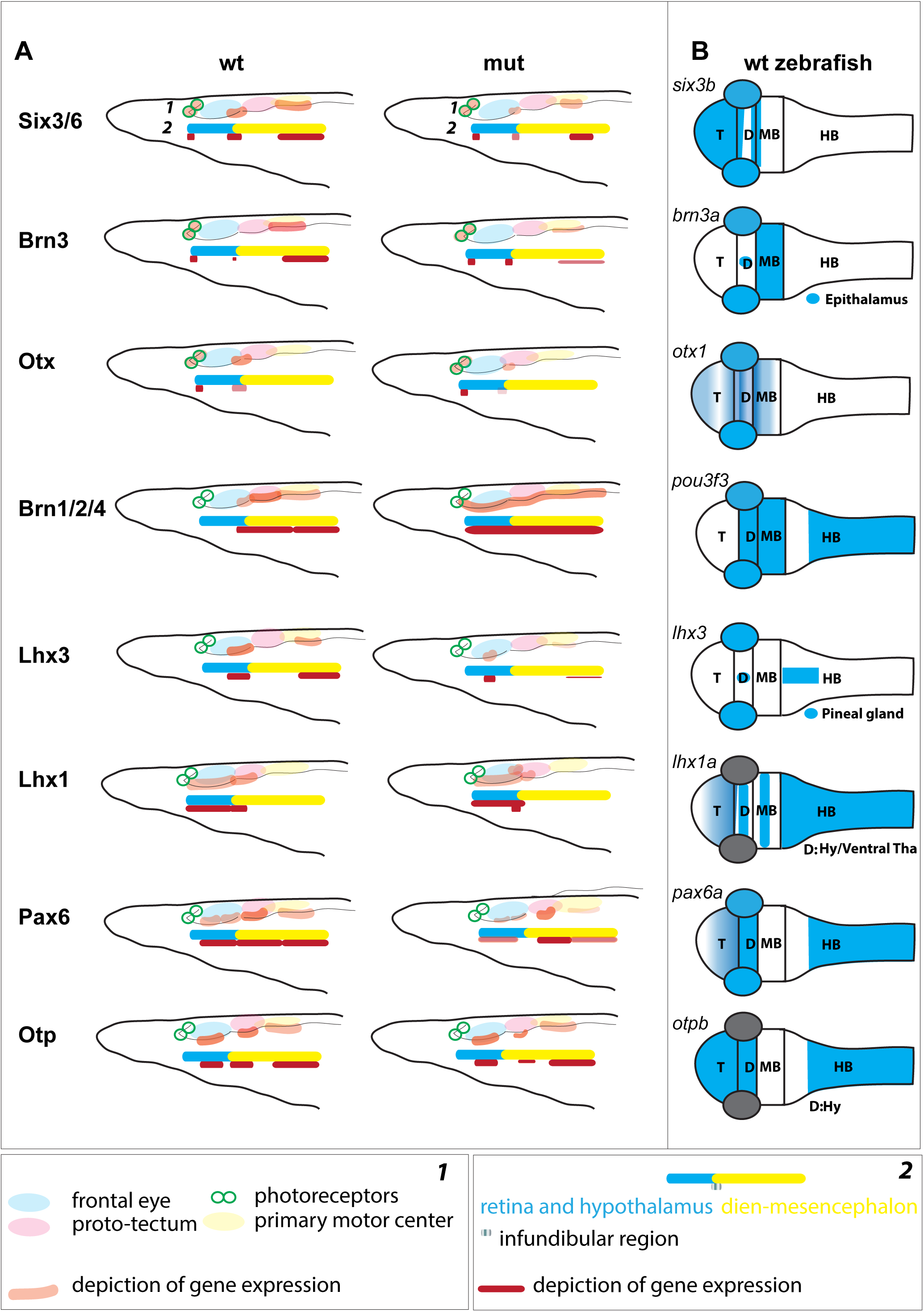
(A) Schematic illustration of gene expression in the anterior central nervous system of amphioxus, which is regionalized according to Pergner *et al*. (schematics ***1***) or Lacalli (schematics ***2***), for both wild type (**wt**) and Pax6 mutants (**mut**). (**B**) Schematic illustration of the orthologous gene expression (*six3b* (*64*), *brn3a* (*65*), *otx1* (*48*), *brn1/2/4 (pou3f3,* http://zfin.org), *lhx3* (*66*), *lhx1a* (*67*), *pax6a* (*48*), *otpb* (*68*)) in the wild type pharyngula stage zebrafish. T – telencephalon; D –Diencephalon; MB – Midbrain; HB – Hindbrain; Hy – Hypothalamus; Tha – Thalamus.

## DISCUSSION

### Composite structure of the chordate brain

It has been argued that the central nervous system of chordates is intricately regionalized, characterized by a complex, gene-specific configuration of the rostral brain as defined by various studies (*28, 29, 31, 32, 35–37*). At least two regions of the brain, the anterior and posterior, are recognized (*31, 32, 38*). These regions are separated from each other by the junction that resembles *zona limitans intrathalamica* (ZLI), a feature molecularly defined in hemichordates and thought to correspond to the infundibular region located at the border between frontal eye and proto-tectum in amphioxus (Fig.5A) (*32, 39*).

In four-day-old larvae of *Branchiostoma floridae*, we observed the presence of *Six3/6* and *Otx* in putative photoreceptors, consistent with previous findings from two-day-old larvae of *Branchiostoma floridae* (*28*) and four-day-old larvae of *Branchiostoma lanceolatum* (*29*). We did not detect *Pax6* in the photoreceptors of our samples, consistent with similar observations in *Branchiostoma lanceolatum* larvae (*29*). However, this contrasts with the weak expression of *Pax6* observed in photoreceptors of two-day-old larvae (*28*), suggesting that the role for *Pax6* is limited to early photoreceptor development. In amphioxus *Pax6* mutants, the anterior frontal eye, including photoreceptors, appeared unaffected. No examinations of possible changes in the photoreceptors have been conducted in *Xenopus* and medaka *Pax6* knockouts (*14, 25, 26*). In *Xenopus Pax6* mutants, the retina is present but disorganized (*25*). In mice, retina-specific *Pax6* gene ablation disrupts normal differentiation program leading to the complete absence of all mature retina neurons (*40*).

A somewhat unexpected finding of our study is the conspicuous expression of *Brn3* in ciliary photoreceptors, an apparent divergence from the situation in vertebrate retinas where *Brn3* is typically absent from photoreceptors and is found in ganglion cells (*41*). A specialized subset of the vertebrate *Brn3*-positive retinal ganglion cells is intrinsically photosensitive due to the expression of the rhabdomeric type opsin (Opn4, melanopsin) (*42, 43*). The presence of *Brn3* in amphioxus ciliary photoreceptors lends support to the hypothesis of a shared ancestral origin between photoreceptors and retina interneurons (*44, 45*).

The notion of homologizing row 2, row 3, and row 4 of the amphioxus frontal eye with the interneuron organization of the vertebrate retina (*29*) apparently warrants further investigation. The previously proposed homology appears less convincing due to the widespread presence of genes in these rows that are also found in other brain regions of vertebrates (Fig.5). For example, we identified *Otp* expression in rows 3 and 4, the proto-tectum, and the primary motor center, a gene not typically expressed in the developing vertebrate eye but a marker of the vertebrate hypothalamus, crucial for neurosecretory cell differentiation (*46, 47*). The presence of *Otp* expression in the frontal eye, which is believed to coincide with both the retina and hypothalamus, aligns with the brain regionalization scheme previously suggested (*31, 32*). Additionally, we discovered *Pax6* expression in the proto-tectum, this contrasts with previous findings (*28, 29*). In vertebrates, *Pax6* shows weak expression in the mesencephalon only during early neurula stages, ceasing at the diencephalon-mesencephalon border in later stages (*2, 14, 25, 26, 31*). However, the expression of *Otp* in the proto-tectum and primary motor center complicates the comparison of this region, called dien-mesencephalon, with the pretectum, thalamus and mesencephalon in vertebrates.

In the border region between the posterior frontal eye and the anterior proto-tectum, which likely corresponds to the infundibular region, we observed elevated expression of *Otx*. In zebrafish, *Otx2* is expressed in the presumptive ZLI region at earlier stages and in the ZLI region at later stages, serving as one of the key factors in establishing this area (*48*). Contrary to the findings of Albuixech-Crespo *et al.* (*31*), some researchers propose that the ZLI is present in the developing brain of amphioxus (*38*) and the infundibular region might represent this (*32*). Our findings lend support to this hypothesis. *Otx*, along with *Wnt8*, *FoxA*, *Hh*, and *Ptch* genes, is expressed in the ZLI-like region of hemichordates (*49*). We were interested in whether the expression of amphioxus *Otx* in the boundary between the posterior frontal eye and the anterior proto-tectum could be observed at earlier stages. We observed the expression of *Otx* and *FoxA* in this region at the tailbud neurula stage (Fig.6Aa-f). Moreover, we detected elevated expression of *Ptch* specifically in this region (Fig.6Ag-i). *Ptch* is a target gene which indicates where Hh signaling is active in amphioxus (*50*). These data suggest that Hh signaling operates in this region similarly to that in vertebrates and hemichordates. Our findings lend support to the hypothesis that a ZLI-like region is present in cephalochordate amphioxus (Fig 6Bc).

**Fig.6.**
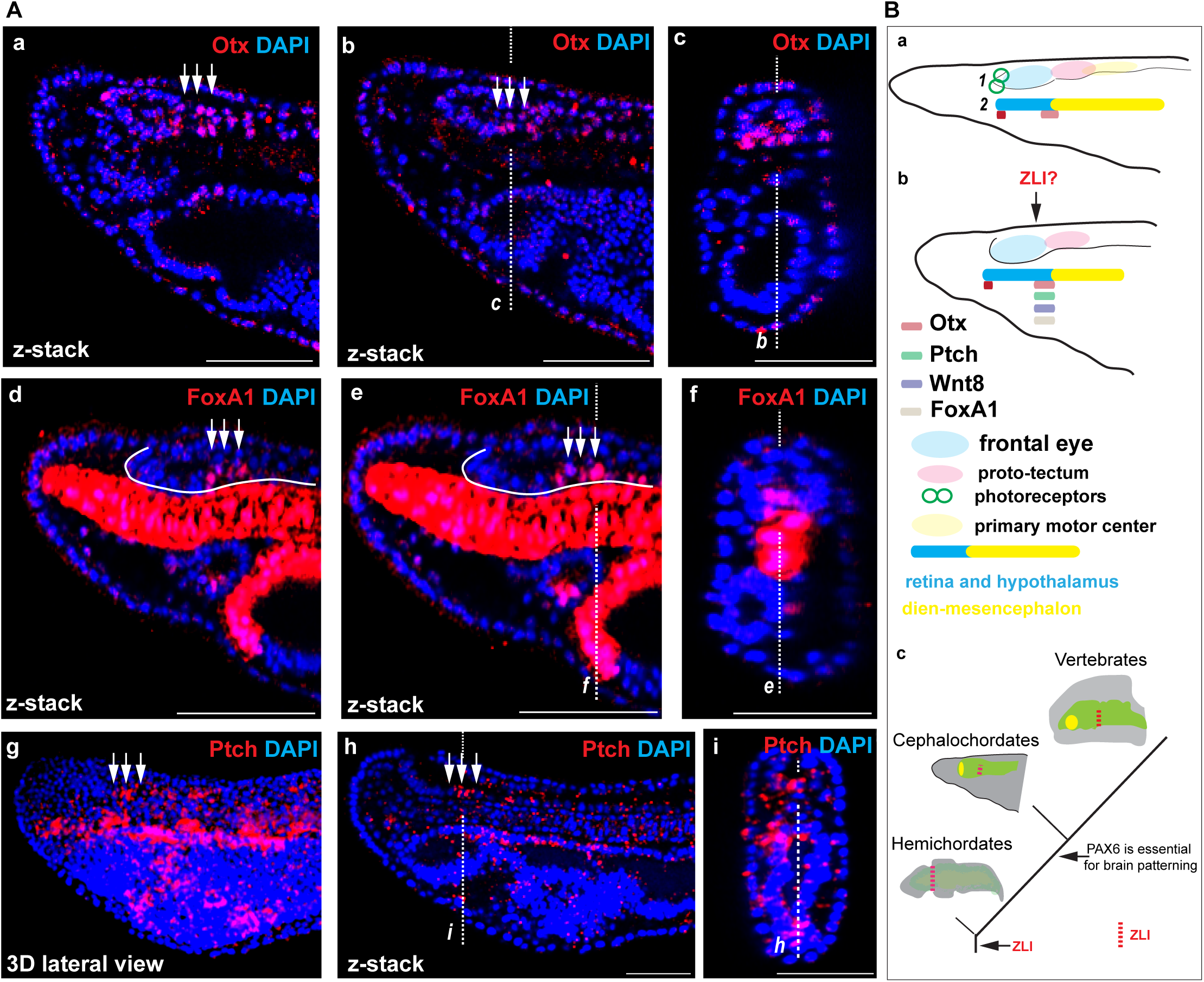
(A) The expression of Otx and FoxA1 proteins (a-c) and *Ptch* mRNA (d-e) in amphioxus tailbud neurula. The positions of individual z-slices (**b-c**, **e-f** and **i-h**) from complete zstacks are indicated with dashed lines (***c-b***, ***f-e*** and ***i-h***). (**B**) Schematic illustration of gene expression in the anterior central nervous system of amphioxus, which is regionalized according to Pergner et al. (schematics ***1***) or Lacalli (schematics ***2***) for *Otx* at the amphioxus four-day-old larvae (**a**) and *Otx*, *Ptch, FoxA,* and *Wnt8* (*69, 70*) at the tailbud neurula (**b**). Proposed scenario of brain patterning in deuterostomes (**c**).

### Conserved role of *Pax6* in the brain regionalization

In our study, we examined amphioxus Pax6 mutants exhibiting a significant decrease in protein activity, apparently due to impaired binding of the paired domain to the DNA-binding site. In the case of heterozygous *Pax6* mutant mice (Sey), which exhibit a noticeable phenotype (*4*), the concentration of functional Pax6 protein is reduced by half. Although not directly comparable, it can generally be stated that in both cases, the regulatory effect of Pax6 transcription factor on gene expression is partially impaired, though not completely abolished. The heterozygous mutation in mice leads to abnormal development of the central nervous system, affecting neuron growth and differentiation in the telencephalon, diencephalon, and metencephalon (*19*). In amphioxus mutants, we observed disorganization of gene expression patterns in various regions of the brain (Fig.5A).

In *Xenopus*, *Pax6* mutant embryos demonstrate changes in the expression of marker genes responsible for telencephalon regionalization, and similar effects have been demonstrated in mice (*51, 52*). It is suggested that *Pax6* plays a crucial role in the regionalization of the telencephalon and diencephalon divisions (*53, 54*). In homozygote *Pax6* mutant mouse embryos, the molecular patterning of the diencephalic regions is compromised, affecting the boundary between the mesencephalon and pretectum, and ZLI (*21, 55*). The molecular markers of the mesencephalon expand into pretectum and the identity of the pretectum is partially shifted towards that of the mesencephalon (*21*). Additionally, the expression markers of the thalamus are downregulated, and genes normally confined to the ZLI are ectopically expressed in the surrounding regions (*20, 55, 56*).

In amphioxus, we observed distinct changes in the expression of individual genes in the posterior frontal eye, proto-tectum and in the primary motor center (the presumed counterparts of the vertebrate diencephalon/eyes, pretectum, and mesencephalon). Most of the examined genes that were expressed in these territories either lost their expression or were significantly downregulated, with the exception of *Brn1/2/4*, which expanded into the anterior region of the frontal eye. Additionally, we observed molecular disorganization in the border region between the frontal eye and the proto-tectum, which is presumed to be the ZLI-like region in amphioxus. Interestingly, the pattern of molecular changes was different from the changes observed in the proto-tectum and primary motor center. For example, Lhx1 expression expanded into the dorsal domain at the anterior border of this region, while its expression was reduced in the proto-tectum. In vertebrates, Lhx1 is expressed in the ZLI, ventral thalamus, and pretectum (*57*). In Pax6 mutant mice, its expression expands into the dorsal thalamus but is reduced in the pretectum (*56*). Combined, these results further support our observation that the border between the frontal eye and the proto-tectum can be recognized as a distinct molecular entity and could thus be homologized to the vertebrate *zona limitans intrathalamica* (ZLI) (*32, 38*).

In summary, our data suggest that the role of *Pax6* gene in the brain patterning is conserved in the chordate lineage and support the hypothesis of the evolutionary continuity of the ZLI-like region in deuterostomes (Fig. 6Bc).

## MATERIALS AND METHODS

### Amphioxus husbandry

Amphioxus *Branchiostoma floridae* adults were housed in seawater at a temperature of 28°C and were fed with algae daily. To induce spawning, the animals were transferred to a temperature of 18°C for at least 6 weeks before being exposed to a heat shock induced by elevating the temperature to 28°C for 24 hours. Following *in vitro* fertilization at room temperature the embryos were raised at a temperature of 25°C.

### Oligonucleotides

Oligonucleotides used for the generation of sgRNA and expression constructs, site-directed mutagenesis, and genotyping are shown in Supplementary Table S1.

### Genome editing

Oligonucleotides zk1770A/B used to make sgRNA constructs were cloned into BsaI site of pDR274(*58*) (pDR274 was a gift from Keith Joung, Addgene plasmid # 42250). Cas9 mRNA was prepared using mMESSAGE mMACHINE T7 ULTRA Kit (Ambion) using plasmid pCS2-nCas9n(*59*) (pCS2-nCas9n was a gift from Wenbiao Chen, Addgene plasmid # 47929). The sgRNAs were transcribed using MEGAshortscript kit (Ambion). A mixture of Cas9 mRNA (100ng/μl) and sgRNA (25ng/μl) was injected into amphioxus eggs, eggs were fertilized, and the developing F0 embryos maintained at 25°C. The adult mature F0 animals were crossed with wild-type animals and the F1 progeny was assayed for mutations by DNA sequencing. Genetic crosses with Pax6ΔQL F1 heterozygotes were used to establish mutant line. Embryos of F2 or F3 generations were used for gene expression analysis. Amphioxus embryos were genotyped using primers zk2059/zk1989QL2/zk614 to distinguish wild type, heterozygotes, and homozygotes, respectively.

### Reporter gene assays

Site-directed mutagenesis of *Branchiostoma floridae* Pax6 cDNA cloned in pKW mammalian expression vector was performed by the Quick-Change kit (Stratagene) using primers zk2027A/zk2027B to generate Pax6ΔQL. The cell culture and transient cell transfection was performed as previously described(*60*). Expression vectors encoding either wild type Pax6 or mutant Pax6ΔQL were co-transfected with Pax6-resposive reporter gene (−350GluLuc (*61*)) and the β-galactosidase expression plasmid serving to normalize the transfection efficiency. Graph and statistical analysis of triplicate biological assays were generated in GraphPad Prism software.

### *In situ* hybridization of amphioxus embryos

*In situ* hybridization followed the protocols described previously (*62*). After being fixed overnight at 4°C with 4% PFA/MOPS solution (0.1M 3-(N-morpholino)propanesulfonic acid, 2mM MgSO4, 1mM EGTA, 0.5M NaCl, pH 7.5), the embryos were transferred to 70% ethanol with DEPC-treated water and stored at −20°C. To generate construct for *Ptch* in situ hybridization probe primers zk1979C/D were used.

The color development was achieved through incubation in Vector blue solution from Vector Laboratories. Images of the embryos were captured using confocal microscopy. The embryos were mounted in glycerol on glass depression slides. Z-stack imaging was conducted using a Leica SP8 confocal microscopes, and analysed with FIJI image analysis software.

### Immunohistochemistry of amphioxus embryos

Antibodies recognizing Pax6, Six3/6, Otx, Brn1/2/4, Brn3, FoxA, Lhx1, and Lhx3 were previously used (*28, 29, 63*). Antibody recognizing amphioxus Otp was prepared as described in Bozzo et al. (*63*). To generate construct for over-expression of Otp protein fragment primers zk1361A/B were used. Embryos for immunohistochemistry were fixed and processed as described in detail before (*29*). The embryos were imaged with Leica SP8 confocal microscope and processed with Fiji ImageJ analysis software. Cells positive for individual markers were counted in wild type and Pax6 mutant embryos. GraphPad Prism software was used to generate individual graphs and analyze statistical significance using Mann-Whitney two tailed test.

## AUTHORS’ CONTRIBUTION

ZK and IK designed the study, performed experiments, and analyzed the data. IK wrote the first draft of the paper. Both authors have approved the final manuscript. The authors declare no conflict of interests.

## ACKNOWLEDGMENTS

We thank Gaspar Jekely for the constructive criticism of an earlier version of the manuscript, Halyna Klymets for technical assistance, Anna Zitova and Veronika Noskova for animal husbandry, Walter Knepel for −350GluLuc, Keith Joung for pDR274 (Addgene plasmid # 42250), and Wenbiao Chen for pCS2-nCas9n (Addgene plasmid # 47929). We acknowledge the Light Microscopy Core Facility, IMG CAS, Prague, Czech Republic, supported by MEYS (LM2018129, CZ.02.1.01/0.0/0.0/18_046/0016045) and RVO: 68378050-KAV-NPUI, for their support with the microscopy presented herein.

---

**Supplementary Table 1.**
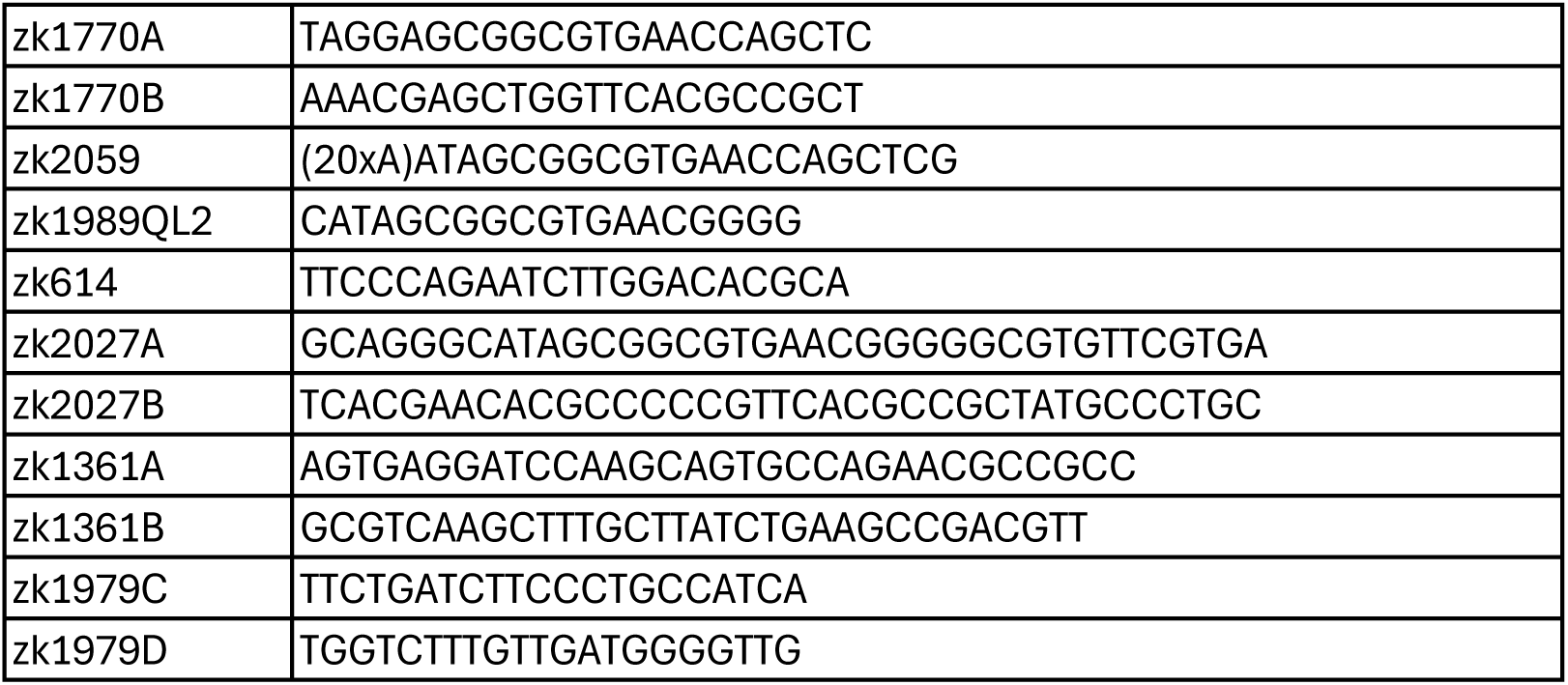

